# From dynamic transcripts to stable proteins: multi-omics revealed regulatory buffering of *Candidatus* Accumulibacter under cyclic conditions

**DOI:** 10.64898/2026.10.10.758232

**Authors:** Ruizhe Pei, Laëtitia Cardona, Pilar Natalia Rodilla Ramírez, Jaspreet Singh Saini, Mathilde Stéphanie Willemin, Maria Pavlou, Christof Holliger

## Abstract

Enhanced biological phosphorus removal (EBPR) relies on polyphosphate-accumulating organisms that experience dynamic fluctuations in redox conditions and substrate availability. How these organisms regulate metabolism across molecular layers under cyclic operation remains incompletely understood. Here, we investigated molecular regulation in a long-term, stable aerobic granular sludge system dominated by *Candidatus Accumulibacter*. We combined metagenomics, time-resolved metatranscriptomics, and matched metaproteomics across three consecutive operational cycles. Genome-resolved metagenomics revealed an *Accumulibacter*-dominated community encoding canonical EBPR functions. Time-resolved metatranscriptomics revealed extensive and reproducible transcriptional dynamics affecting hundreds of genes. These dynamics did not align with discrete EBPR phases or instantaneous environmental transitions, but instead unfolded gradually over the cycle. Despite pronounced transcript-level heterogeneity, aggregation at the functional level revealed strong consolidation, with only a limited number of KEGG orthologs exhibiting coherent temporal regulation. Quantitative metaproteomics revealed a stable proteome across EBPR cycles, with no individual proteins or functional categories showing significant differential abundance despite direct transcript-protein mapping and time-matched sampling. This indicated that transcriptional dynamics did not translate into measurable changes in protein pool size on the time scale of individual cycles, consistent with slow, growth-coupled proteome turnover relative to cycle duration. Together, these results support a hierarchical model of regulation in *Candidatus Accumulibacter*, in which dynamic transcriptional responses are buffered at both the functional and protein levels. This organization provides a plausible basis for stable EBPR performance under predictable cyclic operation and demonstrates how time-resolved multi-omics can be used to resolve regulatory dynamics across molecular layers in engineered microbial ecosystems.

## 1. Introduction

Wastewater treatment is essential for protecting both human and environmental health. To prevent degradation of receiving water bodies, key pollutants such as organic carbon, nitrogen, and phosphorus need to be efficiently removed. Phosphorus in particular can be removed either chemically or biologically. Enhanced Biological Phosphorus Removal (EBPR) is a widely adopted biological strategy that relies on the combination of at least one anaerobic and one aerobic stage (Barnard, 1975; Oehmen et al., 2007). In conventional EBPR, the stages are typically spatially separated and biomass is recirculated (Oehmen et al., 2007). In contrast, aerobic granular sludge (AGS) technology operates in sequencing batch reactors, where treatment phases are separated in time rather than space (Morgenroth et al., 1997; Pronk et al., 2015). AGS operational conditions promote the formation and retention of dense, fast-settling microbial granules, enabling high biomass concentrations, compact reactor design, and simultaneous removal of carbon, nitrogen, and phosphorus (Morgenroth et al., 1997; De Kreuk et al., 2005). Full-scale studies have further shown that AGS can achieve robust nutrient removal with reduced energy demand and reactor footprint compared with conventional activated sludge systems (Pronk et al., 2015). Therefore, AGS has emerged as a compact and energy-efficient platform for biological nutrient removal, including EBPR.

The performance of EBPR systems, both conventional and granular, depends strongly on polyphosphate-accumulating organisms (PAO), among which *Candidatus* Accumulibacter is one of the most extensively studied model lineages (He & McMahon, 2011; Oehmen et al., 2007). Despite its ecological and engineering importance, *Accumulibacter* has not been isolated in pure culture, limiting direct physiological characterization. Consequently, current understanding of its metabolism and regulation under dynamic EBPR conditions relies largely on culture-independent approaches, including metagenomics, metatranscriptomics, metaproteomics, and *in-situ* chemical imaging (Cardona, Ramírez, et al., 2025; Fernando et al., 2019; García Martín et al., 2006; Oyserman et al., 2016; Wilmes, Wexler, et al., 2008).

The first genome-resolved reconstruction of *Accumulibacter* provided a metabolic blueprint for EBPR, revealing the genetic basis for phosphate uptake, polyphosphate turnover, intracellular carbon storage, glycogen cycling, and energy generation (García Martín et al., 2006). Subsequent metatranscriptomic studies showed that core EBPR pathways are broadly maintained across anaerobic-aerobic cycling, while selected metabolic modules, including central carbon metabolism and redox-balancing pathways, respond dynamically to changing environmental conditions (He & McMahon, 2011; Oyserman et al., 2016). More recent work further revealed metabolic and regulatory variability among coexisting *Accumulibacter* lineages, suggesting that stable EBPR performance may emerge from populations with divergent physiological strategies. However, transcript abundance does not necessarily translate into protein abundance or metabolic activity, and metaproteomic studies of EBPR systems remain comparatively limited (Kleikamp et al., 2023; Wilmes, Wexler, et al., 2008; Ziliani et al., 2023). Moreover, transcript and protein pools change on different time scales: proteome remodeling depends largely on growth-associated synthesis and dilution and therefore proceeds slowly in slow-growing organisms, whereas the anaerobic-aerobic transitions of an EBPR cycle occur within minutes to hours. Rapid metabolic switching in PAO may instead be mediated by changes in intracellular storage polymers and metabolite pools, as illustrated by the dynamic shifts in adenylate and uridylate energy charge during anaerobic acetate uptake and polyphosphate degradation in EBPR granular biofilms (Van Ede et al., 2025). Yet it remains unclear how transcriptional dynamics are propagated, buffered, or reorganized at the protein level during cyclic EBPR operation, highlighting the need for time-resolved measurements across multiple molecular layers in stable and well-characterized systems.

In this study, we investigated molecular regulation in a long-term, stable AGS-EBPR reactor dominated by *Accumulibacter.* Genome-resolved metagenomics was used to characterize the microbial community, yielding a curated reference set of 53 metagenome-assembled genomes (MAGs) representing the dominant populations. Within this community, one MAG classified as *Ca.* Accumulibacter necessarius was consistently dominant and accounted for the majority of both metatranscriptomic and metaproteomic signal. Thus, while the metagenomic analysis provides community-level context, subsequent analyses of transcriptional dynamics, differential gene expression, time-series regulation, and proteomic patterns focused on this dominant *Accumulibacter* population. Specifically, we asked whether cycle-scale transcriptional dynamics propagate to the proteome or, consistent with slow proteome turnover, are buffered at the protein level.

## 2. Materials and Methods

### 2.1 Reactor configuration and operation

A bubble-column sequencing batch reactor (D = 6.2 cm, working volume 2.4 L) was operated in fill-draw mode at 18 ± 1 ℃, pH 7.5, using a double-walled jacket with recirculating water. The reactor was operated as described in detail previously (Cardona, Saini, et al., 2025). Briefly, the reactor was inoculated with 5 gMLVSS/L of a homogenized activated sludge collected from the anaerobic tank of the ARA Thunersee wastewater treatment plant (Thun, Switzerland) operating biological phosphorus removal. A synthetic medium was prepared as two separate concentrates containing either the carbon (acetate:propionate) or the nitrogen and phosphate source. The media composition can be found in the Supplementary Information (SI).

During the startup phase, to maintain a COD:P ratio of 12, influent COD was ramped from 50 to 300 mgCOD/L over the first 15 days. Phase durations were varied adaptively during the startup phase of the reactor to ensure complete substrate consumption during the anaerobic phase and biomass retention. Once granules became visible and the reactor reached steady state with the targeted loading, the durations of the phases in each cycle were kept constant. Specifically, each cycle consisted of: (1) 5 minutes N_2_ purge, (2) 60 minutes anaerobic feeding phase (acetate and propionate 1:1 COD ratio) under mixing with bubbling nitrogen gas, (3) 30 minutes anaerobic phase to ensure complete consumption of the carbon substrates, (4) 150 minutes aerobic phase by sparging compressed air, (5) 15 minutes N_2_ purge, (6) 3 minutes settling phase, and (7) 10 minutes decanting step to remove 50% of the supernatant.

### 2.2 Sampling

After 15 days of startup, the reactor was operated under steady-state conditions until day 70, when the samples were taken during three consecutive cycles. In each cycle, biomass and liquid samples were collected at seven time points as shown in Fig. 1, specifically during the feeding phase at 2, 15 and 50 minutes; 2 minutes into the anaerobic phase; and in the aerobic phase at 2, 60 and 150 minutes (i.e., 7, 20, 55, 67, 97, 155 and 245 min after the start of the cycle). At each time point, 14 mL of mixed liquor, 7 mL each from the mid-column and bottom, was withdrawn. The sample was centrifuged at 4200 x g at 4 ℃ for 1 minute. The supernatant was filtered through 0.2 µm syringe membrane filters for chemical analysis and the pellet was divided for metatranscriptomics and metaproteomics.

**Figure 1.**
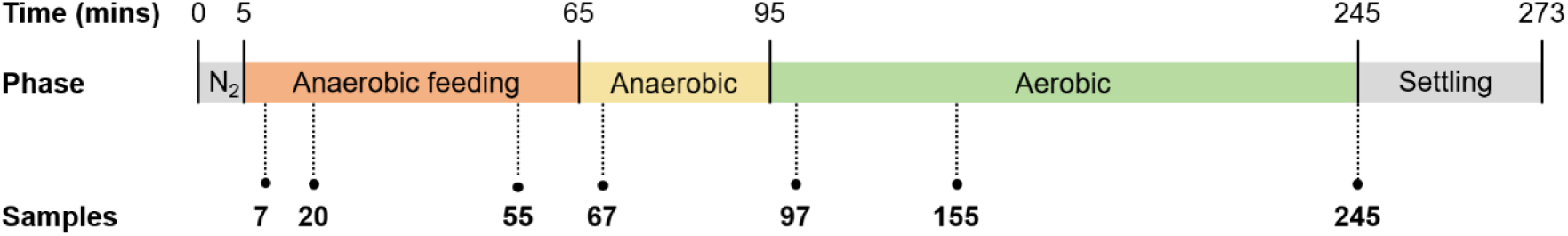
Operational phases and time-resolved sampling scheme.

For metatranscriptomics, the pellet was prepared as described before (Cardona, Ramírez, et al., 2025). The pellet was resuspended in RNAprotect Tissue Reagent (Qiagen, Germany), homogenized, snap-frozen in liquid nitrogen and stored at -80 ℃ until RNA extraction. For metaproteomics, the pellet was resuspended in refrigerated 10 mM HEPES buffer to a final biomass concentration of 0.1 g/L. Homogenization was performed using a glass homogenizer (50 cm^3^, Carl Roth, Germany) with a 0.15-0.25 mm clearance at 700 rpm for 10 up-and-down strokes. A second homogenization was carried out using a Dounce homogenizer (pestle B, Carl Roth, Germany) with 20 manual strokes with rotation. The homogenized biomass was sequentially filtered through 60 µm, 30 µm, and 11 µm filters. The filtrate was aliquoted, snap-frozen in liquid nitrogen, and stored at -80 °C.

Additionally, the biomass taken in the first cycle during the anaerobic phase was collected for metagenomics. The sample was centrifuged at 4200 x g at 4°C for 5 minutes. The pellet was washed twice with 5 mL of phosphate buffer saline (PBS) then centrifuged under the same condition. The washed pellet was resuspended in 3 ml of PBS, homogenized with a glass homogenizer with a clearance between 0.15 and 0.25 mm (50 cm^3^, Carl Roth, Germany) then stored at -20 ℃ until DNA extraction.

### 2.3 Chemical Analysis

COD was measured on the filtrates with the COD Hach cuvette test kits LCK314 and LCK514 and a DR3900 spectrophotometer (Hach Lange, USA). The concentrations of anions (P-PO_4_ ^3-^, N-NO ^-^, and N-NO ^-^) were measured by ion chromatography (IC, ICS-90, IonPacAS14A column) with an electrical conductivity detector (Dionex, Switzerland). In addition to the samples collected for multi-omics analysis, influent and pre-withdrawal samples were collected for nutrient analysis.

### 2.4 Metagenomics

For DNA extraction, 200 μL of homogenized biomass were mixed with 400 μL of elution buffer (10 mM Tris, 0.1 mM EDTA) and 100 μL of lysozyme solution (25 mg/ml). The sample was then incubated at 37 ℃ for 1 hour. Afterwards, DNA was extracted using Maxwell 16 automated DNA purification system following the manufacturer’s instructions (Promega Corporation, Switzerland). The DNA concentration of each DNA extraction was measured with a NanoDrop ND1000 spectrophotometer (Witec AG, Switzerland).

Libraries were prepared from 100 ng of DNA with the Illumina DNA Prep Kit (Illumina, USA) and Unique Dual Indexed oligonucleotides (IDT). PCR amplification was performed for 5 cycles. Final purification was performed twice with 0.6x and 0.54x volume of SPRIselect beads (Beckman Coulter, USA) in order to select for fragments ≥ 600bp. Libraries were quantified by Qubit and their quality assessed on a Fragment Analyzer (Agilent Technologies). Libraries were pooled and circularized with the Element Biosciences Adept Rapid PCR-Free protocol. In order to estimate the loading for the AVITI sequencer, the circularized libraries were quantified by qPCR. This qPCR was done using the Power SYBR green PCR Master Mix in conjunction with the primers provided in the Element Biosciences Adept Rapid PCR-free compatibility kit (qPCR Primer Mix 2, ref: 270-00060). The standard curve was generated using Element Bioscience qPCR Standard 2 (ref: 270-00059). Mix composition and cycling conditions followed the recommendations of the Element Biosciences Adept Rapid PCR-free compatibility kit protocol. The prepared library was sequenced by the Lausanne Genomic Technologies Facility on an Element Biosciences AVITI for 600 cycles (2 x 300 bp). Base calling and demultiplexing were done with bases2fastq (version: 1.6.0), with filter mask R1:N15Y* and --legacy-fastq option for fastq format compatibility with Illumina.

MAGs were reconstructed from quality-controlled and trimmed reads using SPAdes v3.15.5 (Prjibelski et al., 2020) for the assembly and MetaBAT2 v2.12.1 (Kang et al., 2019) and SemiBin2 v2.1.0 (Pan et al., 2023), alongside the competitive approach VAMB v4.1.2 (Nissen et al., 2021). MAGs were dereplicated using dREP v3.5.0 (Olm et al., 2017) and the quality was evaluated using CheckM2 v1.0.1 (Chklovski et al., 2023). GTDB-Tk v2.4.0 (release 220) (Chaumeil et al., 2022) alongside CAT/BAT toolkit v0.6.1 (Von Meijenfeldt et al., 2019) were used for taxonomic assignment. Prodigal v2.6.3 (Hyatt et al., 2010) was used to extract protein-coding sequences and DRAM v1.4.6 (Shaffer et al., 2020) and eggNOG using emapper-2.0.1 and emapper DB: 2.0 (Cantalapiedra et al., 2021) were used to annotate the genes. The complete pipeline is described in the SI.

MAGs with a minimum completeness of 60% and contamination below 5% based on CheckM2 were kept to build the genomics reference database for the metatranscriptomics and metaproteomicsanalyses. To improve functional coverage in downstream analyses, an exception was made for the genus *Contendobacter*. One MAG was manually retained based on a compromise between completeness and contamination (72.61% completeness; 6.86% contamination), resulting in a curated reference set of 53 MAGs.

### 2.5 Metatranscriptomics

For metatranscriptomics, RNA extraction, sequencing, and processing were previously described (Cardona, Ramírez, et al., 2025). Briefly, the RNA was extracted using chloroform, purified with Direct-zol RNA Miniprep (Zymo Research, USA) and DNA was removed using TURBO DNA-free kit (Thermo Fisher Scientific, USA) and further purified using Agencourt RNAClean XP magnetic beads (Beckman Coulter, USA). Afterwards, the ribosomal RNA (rRNA) was removed using QIASeq FastSelect 5S/16S/23S kit (Qiagen, Germany) following the manufacturer’s instructions with 1 µg of total RNA. Libraries were then generated using the TruSeq Stranded mRNA sample preparation kit (Illumina, USA). The libraries were sequenced on NovaSeq 6000 in paired-end mode (2 x 150 bp) at the Lausanne Genomic Technologies Facility, University of Lausanne, Switzerland.

Raw reads were assessed using FastQC v0.11.9 (Andrews, 2010) and summarized using MultiQC v1.13 (Ewels et al., 2016). Adapter removal and quality trimming were performed using BBDuk from BBMap v39.01 (Bushnell, 2014) with the parameters ktrim=r, k=23, mink=11, hdist=1, tpe, tbo for adapter trimming and qtrim=rl, trimq=20, minlen=50, maq=20, maxns=1 for quality filtering. Ribosomal RNA (rRNA) reads were removed using SortMeRNA v4.3.6 (Kopylova et al., 2012), in combination with rRNA databases from SILVA (Quast et al., 2012) and Rfam (Kalvari et al., 2018).

The filtered, non-rRNA reads were then mapped to the MAGs obtained as described in the previous section using Bowtie2 v2.4.1 (Langmead & Salzberg, 2012) with the parameters --very-sensitive, -X 1000, and -k 10. Alignments with less than 95% identity were removed using msamtools (https://github.com/arumugamlab/msamtools) with the filter -p 95 option. Statistics of the mapping results before and after filtering were obtained using SAMtools (Li et al., 2009). Read quantification was performed using featureCounts v2.0.1 (Liao et al., 2014). To account for multi-mapped reads, the parameters -O -M -B --fraction were used, while -O -M -B --primary -Q 2 was applied to obtain counts for uniquely aligned reads.

### 2.6 Metaproteomics

#### Sample preparation

Biomass samples (50 mg) were lysed in 8 M urea supplemented with protease inhibitors and disrupted by sonication (5 x 5 s, 60% amplitude). After centrifugation (13,000 rpm, 15 min, 4 ℃), supernatants were collected, snap-frozen, and stored at -80 ℃. Protein concentrations were determined using Pierce^TM^ BCA Protein Assay Kit (Thermo Fisher Scientific, USA).

Protein extracts were briefly separated by SDS-PAGE (12%), and gel bands containing concentrated proteins were excised. Following washing, proteins were reduced (10 mM dithioerythritol, 1 h, 56 ℃), alkylated (55 mM iodoacetamide, 45 min, dark), and digested overnight at 37 ℃ with trypsin. Peptides were extracted, dried, and desalted using SDB-RPS StageTips.

For quantification, peptides were labelled using TMTpro 18-plex reagents according to the manufacturer’s protocol. Labelling reactions were performed for 90 min at room temperature and quenched with hydroxylamine. Samples were distributed across two TMT sets, each including a pooled reference (“bridge”) sample for cross-set normalization. Labelled peptides were combined at a 1:1 ratio, desalted, and fractionated into 12 fractions using OFF-Gel electrophoresis prior to LC-MS/MS analysis.

#### TMT-based metaproteomics

Peptide fractions were analyzed by nanoLC-MS/MS using an Orbitrap-based mass spectrometer operated in data-dependent acquisition mode. Full MS scans were acquired at high resolution, followed by high energy collision dissociation fragmentation of the most intense precursor ions with dynamic exclusion.

Raw data were processed in Proteome Discoverer (v3.0) using Sequest HT and MSFragger against a database derived from 53 MAGs supplemented with common contaminants. Trypsin specificity was applied with up to two missed cleavages. Carbamidomethylation and TMT tags were set as fixed modifications, and methionine oxidation as variable. Peptide and protein identifications were filtered at 1% FDR, requiring at least two peptides per protein.

Reporter ion intensities were normalized using a multi-step approach implemented in R. First, sample loading normalization was applied within each TMT set. Second, inter-set variation was corrected using internal reference scaling based on bridge samples. Finally, trimmed mean of M-values (TMM) normalization was performed to account for compositional biases. A complete description of the sample preparation and preprocessing is provided in SI.

### 2.7 Statistical Analysis

All statistical analyses were performed using R v4.3.2 and RStudio v2023.12.0+369. Low-count metatranscriptomic features were removed using filterByExpr in edgeR v4.0.16 (Robinson et al., 2010), with a minimum count threshold of 10. A pseudocount of 0.5 was used for zero values. Principal component analysis (PCA) was performed using prcomp after centered log-ratio (CLR) transformation. Differential gene expression (DGE) analysis was performed on the low-count-filtered data using edgeR, DESeq2 v1.42.1 (Love et al., 2014) and limma-voom v3.58.1 (Ritchie et al., 2015). A gene was considered as differentially transcribed if it was detected with at least two of the tools (log2FC > 1.5 and Benjamini-Hochberg-adjusted p < 0.05). The differentially transcribed genes were clustered using pheatmap v1.0.12 (Kolde, 2025) using correlation as distance method and Ward.D2 as clustering method. The profiles of the transcripts were plotted using ggplot2 v3.5.1 (Wickham, 2016) after scaling using Zscore on CLR values. Linear mixed model (lmms, basis p-spline) was performed in addition to the DGE analysis for detecting differentially transcribed genes.

Metaproteomic abundance data were log-transformed and Z-score scaled for visualization. PCA was performed on the log-transformed abundances using prcomp. Differential protein abundance analysis was performed using limma.

## 3. Results and Discussion

Using a time-resolved multi-omics dataset, we examined how a stable EBPR microbial community responds at multiple molecular levels to cyclic changes in substrate availability, redox conditions, and phosphate concentration. We first established process stability and community composition, then resolved transcriptional regulation across consecutive EBPR cycles using complementary differential expression and time-series analyses, and finally assessed how these transcriptional signals were buffered at functional and protein levels. This structure allowed us to evaluate how regulation propagated across molecular layers in an *Accumulibacter*-dominated system.

### 3.1 Stable Reactor Performance and *Accumulibacter*-dominated EBPR Community

The bubble-column reactor was operated under cyclic EBPR conditions for 70 days prior to sampling, allowing the microbial community to reach a stable, EBPR-adapted state (Figs. S1 and S2). Reactor performance demonstrated strong selective pressure for PAO, consistent with previous AGS-EBPR studies showing that anaerobic-aerobic operation with acetate and propionate as the sole carbon sources selects for PAO-associated granular communities and links nutrient removal performance to bacterial population dynamics (Weissbrodt et al., 2014). Amplicon sequencing further confirmed *Accumulibacter* became the dominant genus from ∼day 40 and continued to increase until sampling (Fig. S2).

Time-resolved sampling was conducted across three consecutive EBPR cycles. As shown in Fig. 2, acetate and propionate were effectively consumed during the anaerobic phase, with no measurable residual carbon detected at the onset of aeration in cycles 1 and 3, and only trace concentrations in cycle 2 (acetate: 0.63 mg/L; propionate: 0.56 mg/L). Phosphate removal occurred during the aerobic phase, with effluent concentrations reaching an average of 25.7 ± 23.1 mg P-PO_4_ ^3-^/L by the end of each cycle, with cycle-to-cycle variability.

**Figure 2.**
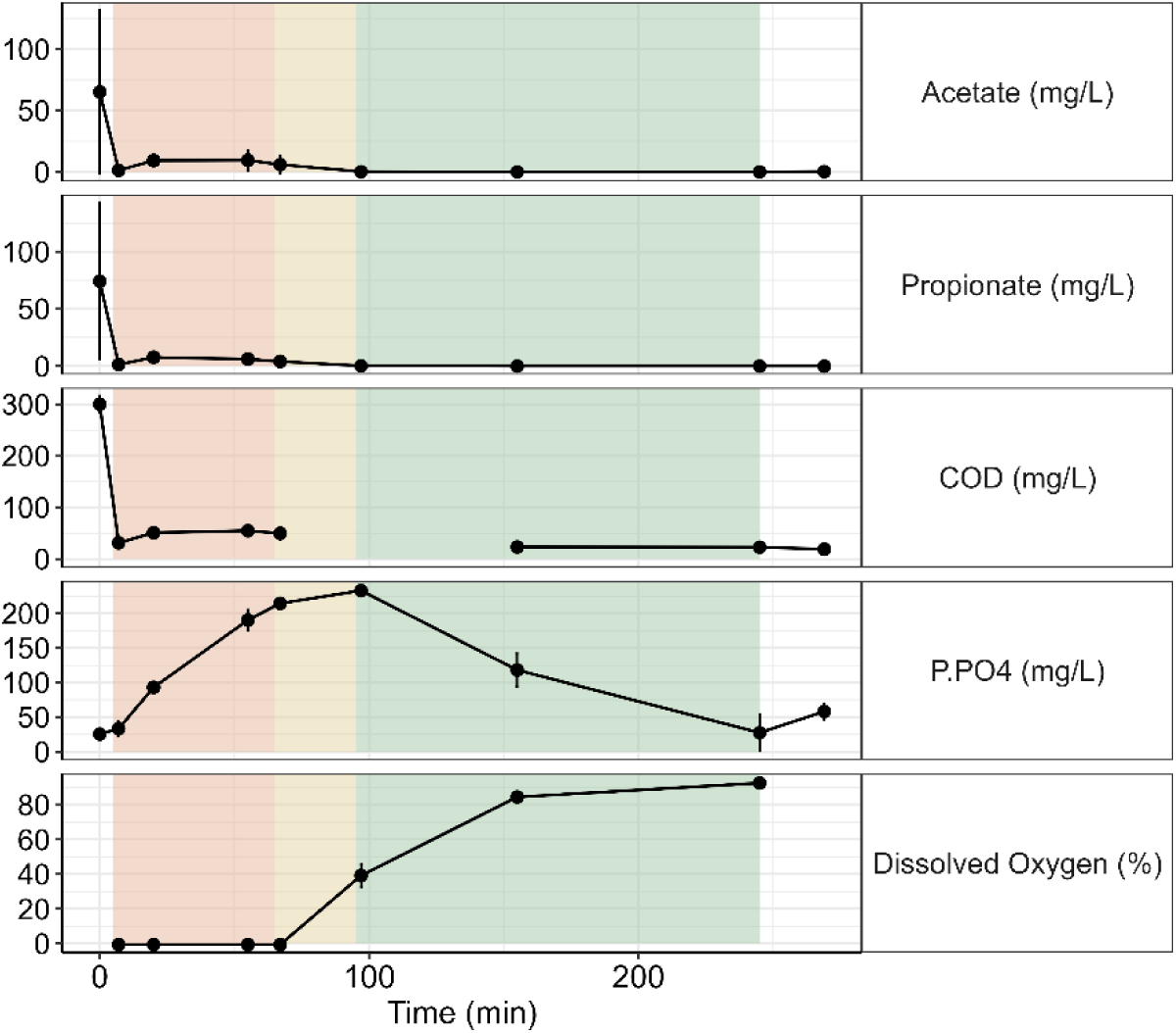
Average reactor performance across the three EBPR cycles used for time-resolved sampling. Points represent mean values across the three sampled cycles and error bars indicate standard deviation. Shaded regions indicate the successive operational phases of the cycle.

Considering the stability of the microbial community and the slow growth characteristics of microorganisms in the AGS system, metagenomic sequencing was performed only at the beginning of the multi-omics sampling campaign. Genome-resolved metagenomic analysis yielded 1126 initial bins, which were refined and filtered to 52 medium-quality MAGs (>60% completeness, <5% contamination). Taxonomic classification with GTDB-Tk revealed that *Accumulibacter* dominated the community. Five MAGs were assigned to this lineage (average completeness: 89.3%; average contamination: 3.2%), collectively accounting for 56.27% of the mapped reads. Notably, a single MAG classified as *Accumulibacter necessarius* represented most of the population and accounted for 53.34% of the total mapped reads. This strong dominance is consistent with previous laboratory-scale EBPR studies in which *Accumulibacter*-enriched communities were used to resolve the genomic and transcriptional basis of EBPR metabolism (García Martín et al., 2006; He & McMahon, 2011; McDaniel et al., 2021; Weissbrodt et al., 2014). Functional annotation further confirmed that the dominant *Accumulibacter* MAGs encoded canonical EBPR pathways, including volatile fatty acid uptake, intracellular carbon storage, polyphosphate metabolism, and respiratory energy generation, consistent with conserved metabolic features reported across *Accumulibacter* genomes (Flowers et al., 2013; García Martín et al., 2006; Petriglieri et al., 2022).

One *Contendobacter* genome that did not initially pass the quality filter (72.61% completeness; 6.86% contamination) was manually retained because members of the *Candidatus* Competibacter/Contendobacter lineage are glycogen-accumulating organisms (GAOs), which are widely detected in EBPR systems and can compete with PAO for volatile fatty acids under anaerobic-aerobic cycling (McIlroy et al., 2014; Oehmen et al., 2006). This resulted in a curated 53-MAG reference set for the downstream analysis.

### 3.2 *Accumulibacter* Transcriptional Dynamics are Gradual and Buffered at the Functional-annotation Level

To resolve how the dominant PAO population responded to cyclic EBPR operation, metatranscriptomes were analyzed across three consecutive reactor cycles. After quality trimming and filtering, approximately 32 to 56.6 million high-confidence mRNA reads were retained per sample, with comparable read loss across samples and no apparent phase-or cycle-specific processing bias (Fig. S3). Across the 53-MAG reference database, metatranscriptomic reads mapped to 132608 of 153818 predicted genes, of which 30448 passed low-abundance filtering for statistical analysis. The transcriptome was strongly dominated by *Accumulibacter*: 87% of retained reads mapped to the five *Accumulibacter* MAGs, and *Accumulibacter necessarius* accounted for 91% of all *Accumulibacter*-mapped reads (Fig. 3). For this MAG, the number of detected transcribed genes reached a plateau despite variation in total mapped read counts, indicating that sequencing depth was sufficient to capture the active transcriptional repertoire of the dominant population (Fig. 3).

**Figure 3.**
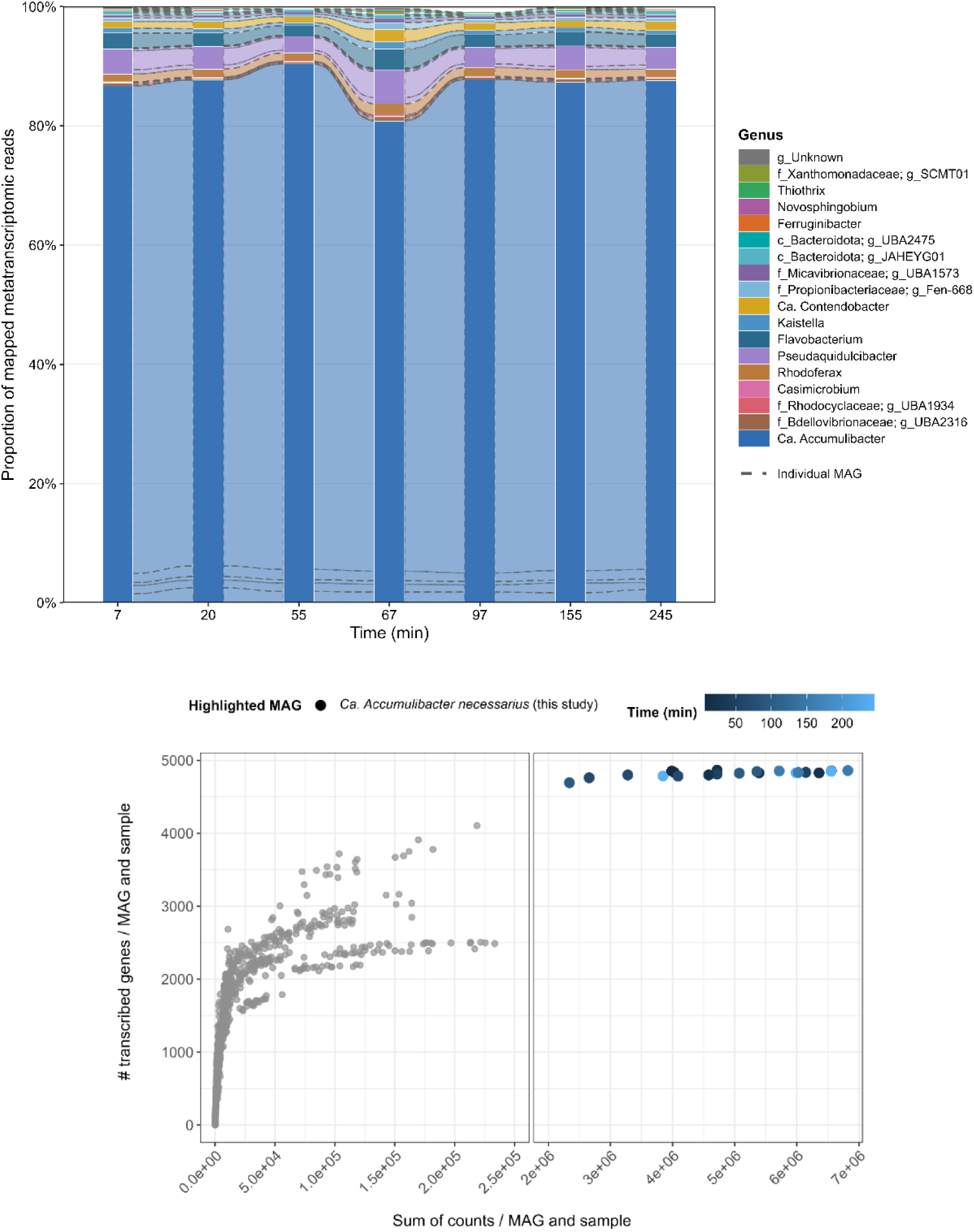
Metatranscriptomic dominance and transcriptional depth of *Accumulibacter*. **Top:** Relative contribution of individual MAGs to the mapped metatranscriptomic reads across the seven sampling times. Colors indicate genus-level affiliation, and dashed lines separate individual MAG trajectories within genera. **Bottom:** Relationship between total mapped transcript counts and the number of detected transcribed genes for each MAG-sample combination. All MAGs are shown in gray, while the highlighted *Accumulibacter necessarius* MAG is colored by sampling time.

Principal component analysis (PCA) was used to assess the global structure of the metatranscriptome. PCA based on all expressed transcripts across the 53 MAGs and PCA restricted to *Accumulibacter necessarius* transcripts produced similar ordination patterns, consistent with the strong contribution of this population to the community-level signal. In both analyses, samples did not separate into discrete clusters corresponding to feeding, anaerobic, or aerobic phases. Instead, samples occupied overlapping regions of ordination space and followed gradual trajectories across the cycle (Fig. 4). This indicates that cyclic EBPR operation did not impose abrupt, phase-separated transcriptomic states at the global level, but rather generated continuous transcriptional reorganization. Previous *Accumulibacter* expression studies similarly showed that many core EBPR functions remain broadly expressed across dynamic anaerobic-aerobic operation, while selected metabolic modules respond to changing acetate, phosphate, oxygen, and intracellular storage conditions (He et al., 2010; Oyserman et al., 2016). Our data extend this view by showing that, in a highly enriched AGS-EBPR system, the dominant *Accumulibacter* population followed a gradual transcriptional trajectory rather than switching between discrete phase-specific states.

**Figure 4.**
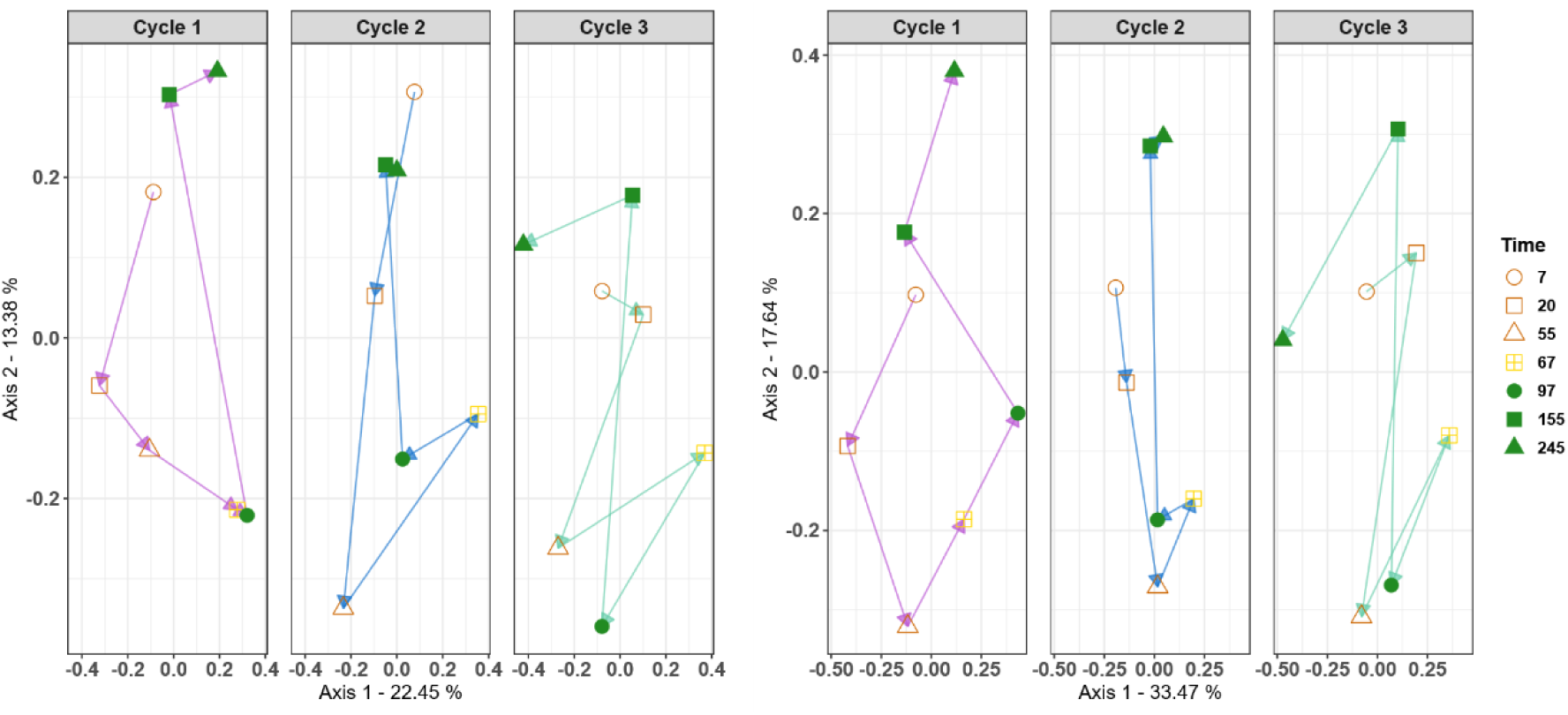
Principal component analysis of metatranscriptomic profiles across three consecutive EBPR cycles. **Left:** PCA based on all expressed transcripts across the 53 MAGs. **Right:** PCA restricted to transcripts assigned to *Accumulibacter necessarius*.

Because of the dominance and transcriptional saturation of *Accumulibacter necessarius*, DGE analysis was focused on this single *Accumulibacter necessarius* MAG. In total, 4796 robustly expressed transcripts were retained. Pairwise DGE analysis was performed using low-count-filtered raw counts, with the three consecutive EBPR cycles treated as replicate cycle observations for comparisons between corresponding time points. To ensure a robust consensus, DGE was assessed using edgeR, limma-voom, and DESeq2, and only genes identified by at least two of the three methods were retained.

Across all contrasts, 386 genes were identified as differentially expressed. The number of differentially transcribed genes varied strongly among contrasts and generally increased with the temporal distance between compared samples (Table 1). Comparisons spanning short intervals early in the EBPR cycle yielded few or no differentially transcribed genes. For example, the earliest contrast (20-7 min) yielded zero differentially transcribed genes across all three statistical methods. Contrasts within the feeding phase identified modest numbers of regulated genes, with 45 and 66 genes detected in the 55-20 and 55-7 contrasts, respectively. Notably, adjacent contrasts across the anaerobic-aerobic redox transition yielded very few differentially transcribed genes (e.g., only 2 genes in the 97-67 comparison), despite the large physiological shift in electron acceptor availability. In contrast, comparisons spanning longer intervals consistently yielded substantially larger gene sets. The contrast between 67 and 7 min identified 135 genes common to at least two methods, while contrasts across extended aerobic operation showed the strongest transcriptional divergence, including 125 genes in the 155-97 comparison, 163 genes in the 245-97 comparison, and 259 genes in the longest contrast tested, 245-67. This pattern indicates that transcriptional divergence accumulates progressively over time rather than occurring as an instantaneous response to environmental transitions.

**Table 1.** Differential gene expression across pairwise time-point contrasts for *Accumulibacter necessarius*. Numbers indicate transcripts identified as differentially expressed by edgeR, limma-voom, and DESeq2 for each contrast. The Common row reports the consensus set retained when a transcript was identified as differentially expressed by at least two of the three methods.

| Contrast (mins) | 20-7 | 55-20 | 55-7 | 67-55 | 67-7 | 97-67 | 155-97 | 245-155 | 245-97 | 245-67 |
| --- | --- | --- | --- | --- | --- | --- | --- | --- | --- | --- |
| edgeR | 0 | 47 | 66 | 37 | 138 | 2 | 129 | 37 | 169 | 269 |
| Limma-voom | 0 | 33 | 65 | 0 | 92 | 2 | 116 | 17 | 138 | 176 |
| DESeq2 | 0 | 52 | 72 | 62 | 172 | 18 | 132 | 23 | 185 | 338 |
| **Common\*** | **0** | **45** | **66** | **37** | **135** | **2** | **125** | **36** | **163** | **259** |

Hierarchical clustering of the 386 differentially expressed genes identified five co-expression clusters with distinct temporal profiles (Fig. S4). Across clusters, expression changes unfolded over tens to hundreds of minutes, further supporting a gradual regulatory regime rather than discrete phase-specific switching. For functional interpretation and visualization, analyses were focused on the subset of 218 annotated genes possessing known or putative functions. The expression profile for the annotated genes is shown in Fig. 5. Cluster 1 was characterized by relatively low expression at the beginning of the cycle, followed by a rapid increase during feeding and early anaerobic operation, peaking near the anaerobic-aerobic transition before progressively declining. Cluster 2 displayed elevated expression early in the feeding phase, followed by a steady decrease through anaerobic operation to reach a minimum early in the aerobic phase, before gradually recovering. Cluster 3 exhibited a contrasting profile to Cluster 1, with reduced expression during the feeding and early anaerobic phases, followed by pronounced induction during aerobic operation to reach maximal expression at later aerobic time points. Cluster 4 showed an early-cycle peak during feeding, followed by a steep decrease that reached its minimum mid-way through the aerobic phase, with partial recovery thereafter. Finally, Cluster 5 was characterized by moderate expression during feeding followed by severe repression during anaerobic operation, before exhibiting a delayed and sustained increase during the aerobic phase, with expression rising gradually rather than abruptly after oxygen exposure.

**Figure 5.**
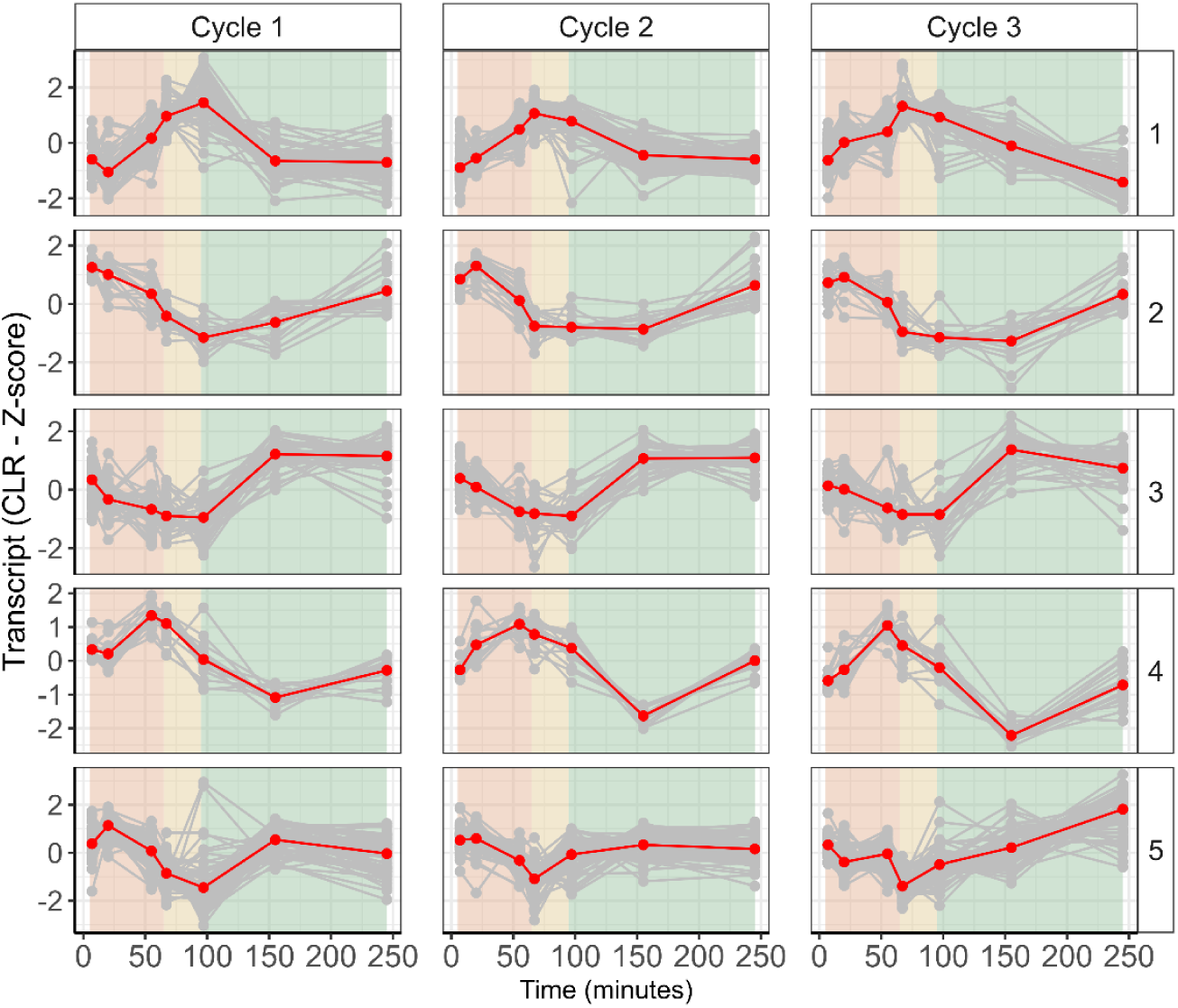
Temporal expression patterns of annotated differentially expressed genes in *Accumulibacter necessarius* across three consecutive EBPR cycles. The 218 annotated genes were grouped into five expression clusters based on their temporal profiles.

Mapping EBPR-relevant functions onto these clusters revealed a coherent functional organization of transcriptional regulation. Genes associated with polyhydroxyalkanoate (PHA) synthesis (*phaC*) and central carbon metabolism (*gltA*, *sdh*, *sucC*) mapped predominantly to Cluster 1, consistent with a transcriptional induction to support the energy demands of early-cycle substrate assimilation. In contrast, genes associated with phosphate transport and polyphosphate-related functions were predominantly assigned to Clusters 3 and 5, aligning with the requirement for polyphosphate regeneration under aerobic conditions.

Beyond canonical EBPR pathways, this continuous regulatory regime also structured a clear temporal partitioning of nitrogen metabolism. Genes encoding the oxygen-sensitive nitrogenase complex (*nifB*, *nifE*, *nifN*, *nifX*, and Mo-Fe subunits), together with the associated Rnf complex, were almost exclusively assigned to Clusters 2 and 4, indicating preferential expression during the anaerobic phase. This pattern is consistent with earlier genome-resolved evidence that *Accumulibacter* encodes nitrogen-fixation-associated genes, but transcript abundance alone does not demonstrate active nitrogen fixation (García Martín et al., 2006). Upon transition to aerobic conditions, *nif* expression was rapidly repressed, coinciding with a shift toward nitrogen scavenging expression. This transition was marked by strong aerobic induction of ammonium and urea transporters (Cluster 3), followed by delayed activation of *urease* (Cluster 5). The data also suggested an anticipatory regulatory strategy for respiration. Multiple subunits of the high-affinity cbb3-type cytochrome c oxidase and ATP synthase clustered in Cluster 1, with transcription increasing during the anaerobic phase and peaking at the onset of aeration. This pre-expression may reflect anticipatory regulation of the respiratory machinery, enabling immediate oxygen utilization and rapid initiation of oxidative phosphorylation upon oxygen availability.

A complementary time-series model (LMMS) supported this interpretation, identifying additional genes with gradual or transient temporal regulation, including EBPR-related genes also detected by pairwise differential expression analysis. We next evaluated the transcripts associated with the same KEGG Orthology (KO). Different copies of the same genes exhibited distinct temporal trajectories. Multiple transcripts annotated to enzymes involved in PHA metabolism (e.g. *phaA*, *phaC*), glycogen turnover (e.g. *glgB*, *glgX*), or phosphate transport and polyphosphate metabolism (e.g. *pitA*, *ppk2*, *ppx*, *pstA*, *pstC*) showed divergent regulation, suggesting selective modulation of paralogs (Fig. 6). However, when transcript abundances were aggregated at the KO level, only 33 KOs showed significant temporal regulation across cycles. This reduction from hundreds of regulated transcripts to a smaller set of regulated functional annotations suggests that copy-specific transcriptional dynamics are partially consolidated at the pathway-annotation level. This interpretation is consistent with genome-resolved metatranscriptomic work showing that coexisting *Accumulibacter* lineages can differ in metabolic potential and regulatory strategy, while maintaining conserved EBPR functions at the population level (McDaniel et al., 2021; Petriglieri et al., 2022).

**Figure 6.**
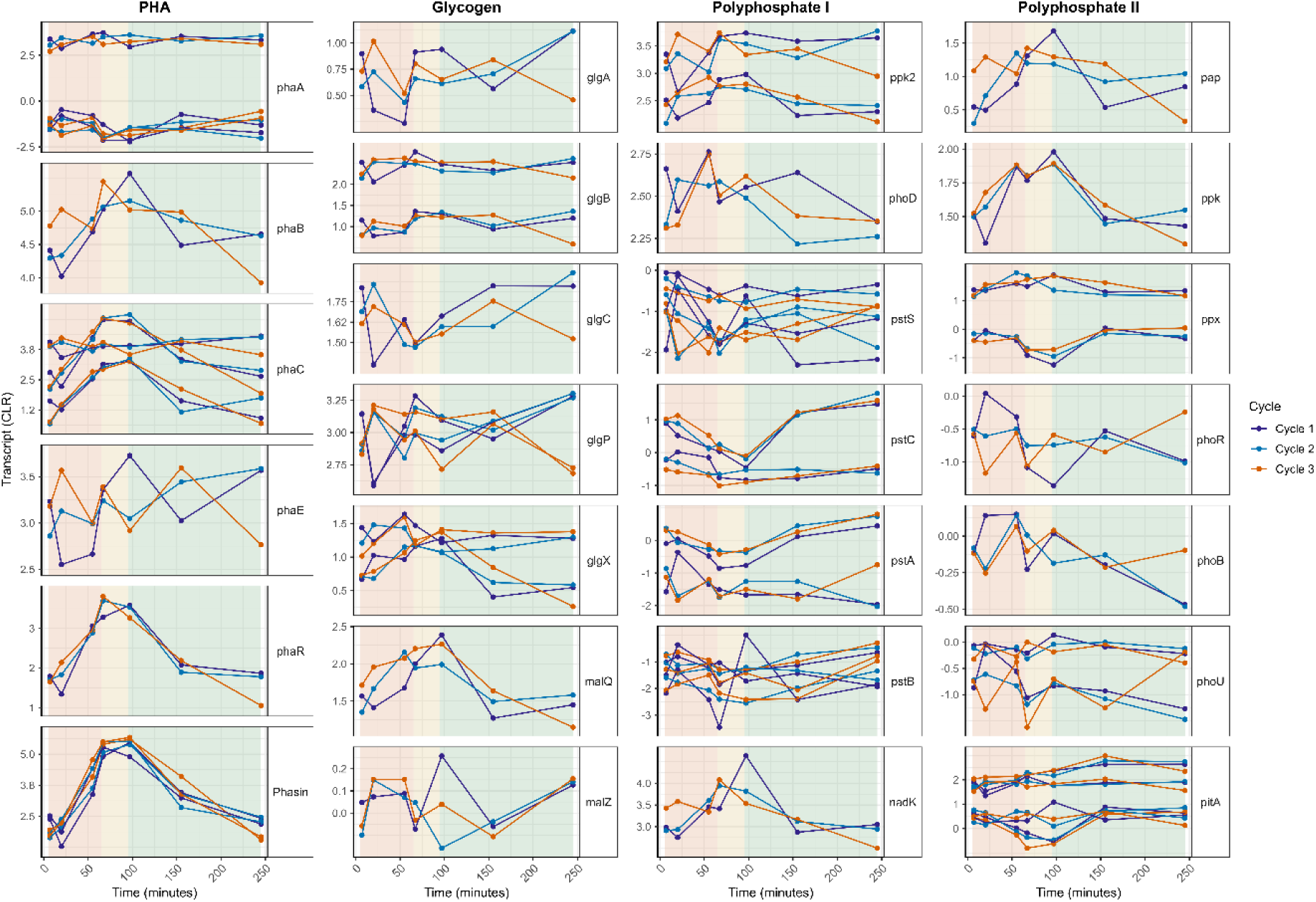
Copy-specific temporal regulation of genes involved in key EBPR functions in *Accumulibacter necessarius*. Temporal expression profiles are shown for transcripts assigned to PHA metabolism, glycogen turnover, polyphosphate metabolism and phosphate transport across three consecutive EBPR cycles.

Together, these results show that transcriptional regulation in *Accumulibacter necessarius* is dynamic but not phase-discrete. Regulatory changes unfolded gradually across the EBPR cycle, with strong gene-and copy-level variability that was partially consolidated when viewed at the functional-annotation level. However, transcript-level consolidation does not necessarily imply equivalent protein-level regulation or metabolic activity. We therefore also examined the metaproteome to determine whether these transcriptional dynamics were propagated, buffered, or reorganized at the protein level.

### 3.3 *Accumulibacter* Protein Expression Remains Stable Across EBPR Cycles

Metaproteomic profiles were generated from the same time points and three consecutive EBPR cycles analyzed by metatranscriptomics. Samples were processed using TMT labeling to improve relative quantitative comparability across time points and reduce technical variation. Protein extraction, labeling, and mass spectrometry acquisition were randomized to minimize run-order bias. Peptide and protein identification were performed against the amino acid sequences of the same curated 53-MAG reference database used for metatranscriptomic analysis, enabling direct comparison between transcript and protein abundance patterns.

Across all samples, 5029 protein groups were identified, of which 3629 were assigned to *Accumulibacter*. These protein groups corresponded to 12330 genes encoded by the five *Accumulibacter* MAGs. For the dominant *Accumulibacter necessarius* MAG, 2340 protein groups were identified and linked to 4714 genes retained in the low-count-filtered metatranscriptomic dataset, enabling direct comparison between transcript and protein abundance patterns. This depth of protein identification indicates that the metaproteome captured a substantial fraction of the detectable protein repertoire of the dominant PAO population. The protein identification level is also comparable to recent metaproteomic studies of AGS and EBPR-associated microbial communities and extends earlier EBPR proteogenomic work that identified hundreds of *Accumulibacter*-associated proteins (Elahinik et al., 2022, 2023; Kleikamp et al., 2023; Van Olst et al., 2025; Wilmes, Andersson, et al., 2008).

Community-level metaproteome structure was explored with PCA based on log-normalized protein abundances (Fig. 7). In contrast to the metatranscriptomic data, metaproteomic profiles showed less clear separation by EBPR phase or cycle. Samples largely overlapped in ordination space, and within-cycle trajectories were weak and inconsistent. Performing PCA on all detected proteins or restricting the analysis to *Accumulibacter necessarius*-derived proteins yielded comparable patterns, although *Accumulibacter necessarius*-only analyses explained a slightly higher proportion of variance. This indicates that the dominant PAO population contributes disproportionately to the detectable proteomic signal, yet even within this population no discrete or reproducible proteomic states emerged.

**Figure 7.**
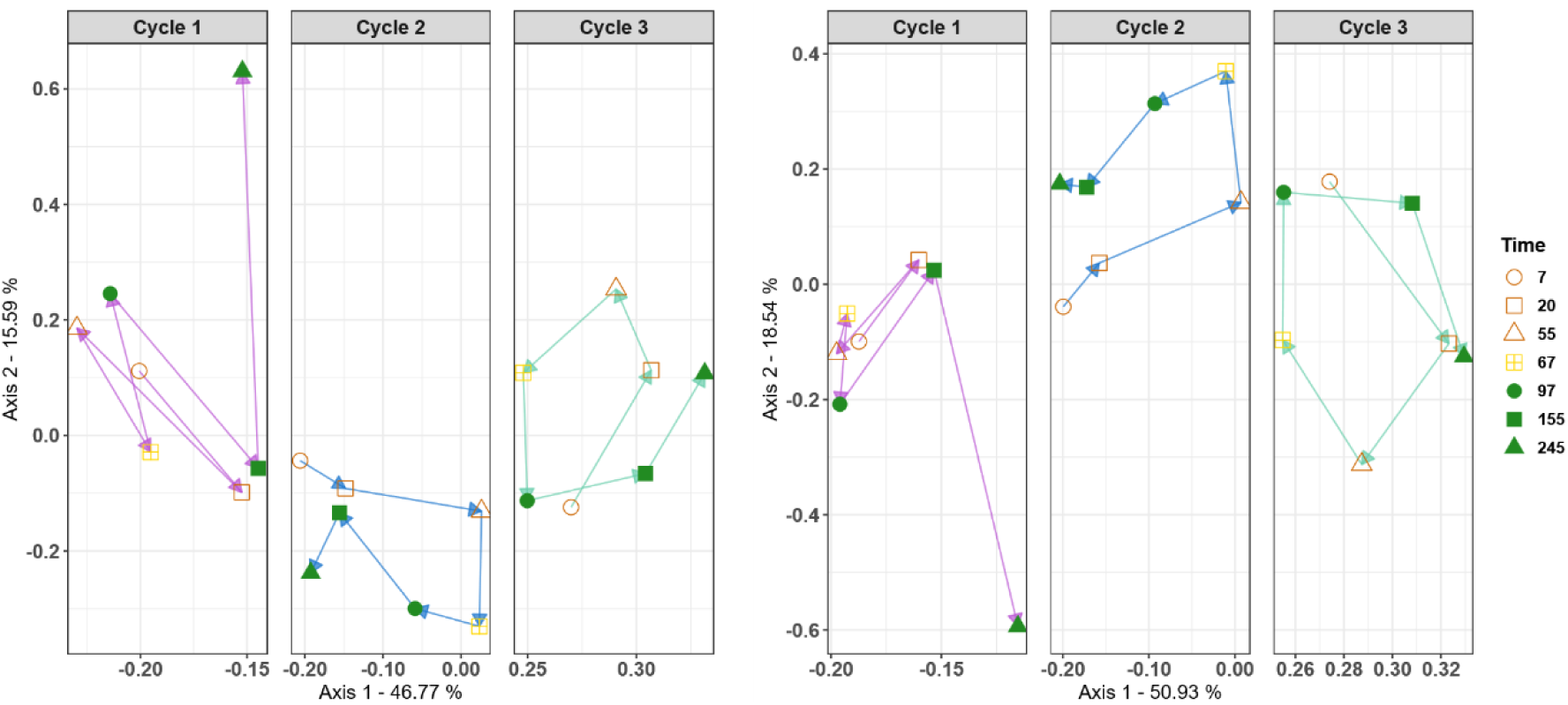
Principal component analysis of metaproteomic profiles across three consecutive EBPR cycles. **Left:** PCA based on all detected proteins across the 53 MAGs. **Right:** PCA restricted to proteins assigned to *Accumulibacter necessarius*.

Consistent with the PCA results, pairwise differential protein abundance analysis identified no significant differences between EBPR phases or time points under conservative statistical thresholds, in contrast to the more pronounced regulation observed at the transcript level. To illustrate this discrepancy, we compared DGE transcripts associated with core EBPR processes with their matched protein profiles shown in Fig. 8. For PHA and polyphosphate metabolism, cycle-resolved transcript changes were often only weakly reflected, temporally shifted, or absent at the protein level. These results indicate decoupling between transcriptional regulation and protein abundance over the time scale of individual EBPR cycles.

**Figure 8.**
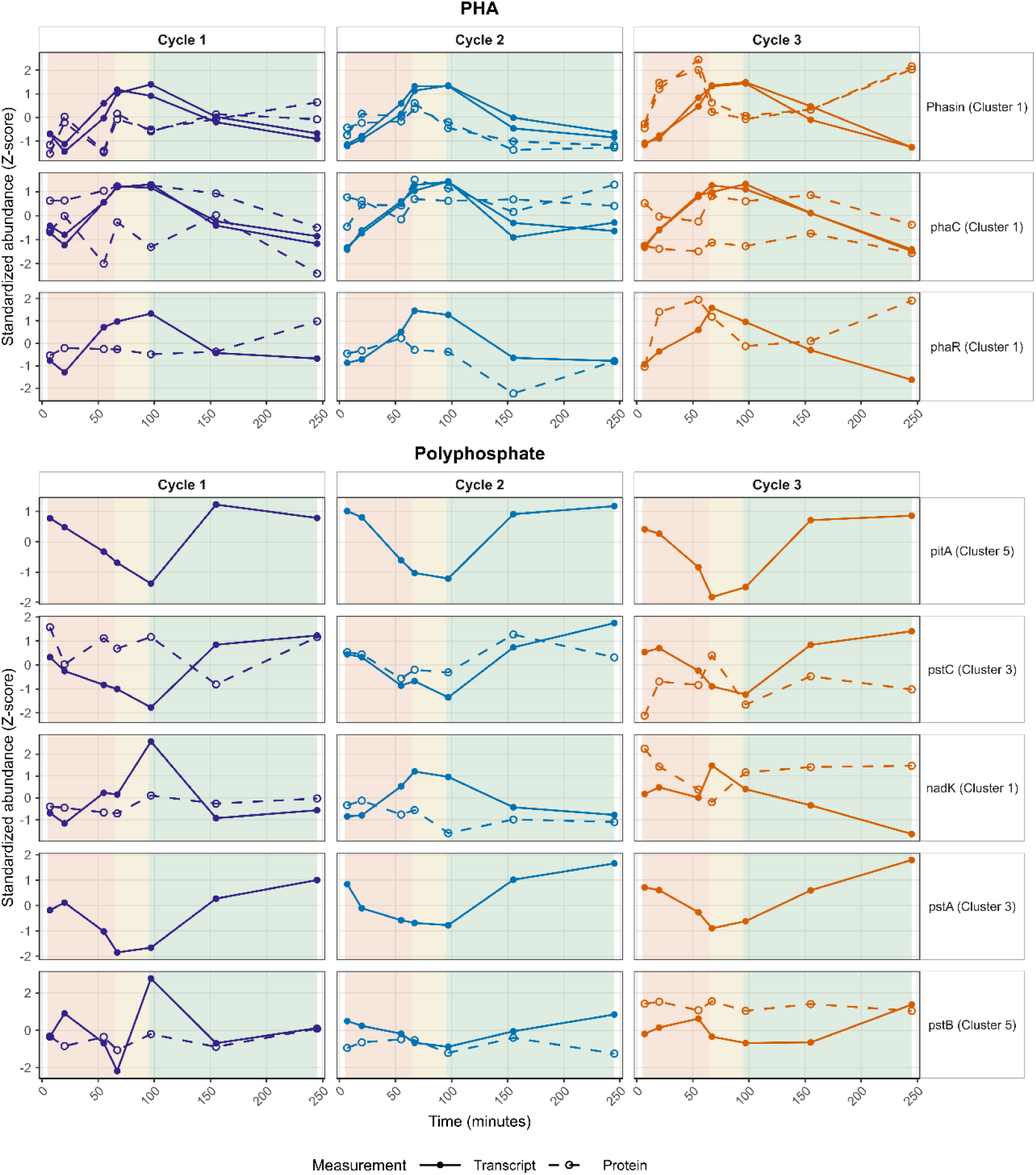
Comparison of transcript and protein abundance profiles for selected differentially expressed genes associated with core EBPR processes across three consecutive cycles. Standardized transcript and matched protein abundances are shown over time, with solid lines representing transcript profiles and dashed lines representing protein profiles.

Aggregating both metatranscriptomic and metaproteomic data at the KO level further reinforced this pattern. While transcript-level aggregation substantially reduced apparent regulatory complexity, revealing a small number of functionally regulated KOs, no corresponding changes were detected at the protein level, either for individual protein groups or after functional aggregation. Thus, even transcriptional regulation that was coherent at the functional level did not translate into measurable changes in protein pool size on the time scale of individual EBPR cycles.

Together, these results indicate that *Accumulibacter* maintains a largely stable proteome across EBPR phases and consecutive cycles, despite more structured transcriptional regulation at the gene and copy levels. This indicates strong buffering at the protein layer, such that transcriptional variability modulates regulatory potential without inducing net changes in protein abundance under routine cyclic operation.

### 3.4 Hierarchical Regulation Buffers Functional Output under Cyclic EBPR Operation

Integrating metagenomic, metatranscriptomic, and metaproteomic data revealed the hierarchical organization of regulation in *Accumulibacter necessarius* under cyclic EBPR conditions. At the transcript level, hundreds of genes exhibited reproducible temporal dynamics across consecutive cycles. These dynamics were gradual and accumulated over time rather than occurring as instantaneous responses to environmental transitions. However, aggregation of transcripts by KO revealed that only a small subset of functions exhibited coherent temporal regulation, indicating consolidation of gene-level variability into a more stable functional output. In contrast to transcriptional variability, protein abundances remained stable across EBPR cycles. Despite time-matched sampling, quantitative TMT-based metaproteomics, and direct mapping between genes and proteins, no individual proteins or functional categories exhibited significant differential abundance. This indicates that transcriptional dynamics do not translate into measurable changes in protein pool size on the time scale of individual EBPR cycles. The solids retention time in the reactor was approximately 21 days, which was more than 100 times the length of a single ∼4.5-h cycle. Under such conditions, rapid or large-scale remodeling of the proteome during individual EBPR cycles is likely neither necessary nor energetically favorable. Instead, *Accumulibacter* appears to maintain a constant proteome sufficient to support EBPR metabolism.

Stable protein abundance does not imply static physiology. Protein synthesis and degradation may occur without net changes in abundance, while post-translational modifications, substrate availability or allosteric regulation may alter enzyme activity without changing total protein levels. In EBPR, rapid metabolic switching is closely tied to the turnover of intracellular storage polymers and energy carriers. Recent nucleotide metabolomics of an EBPR granular biofilm showed dynamic shifts in adenylate and uridylate energy charge during anaerobic acetate uptake and polyphosphate degradation (Van Ede et al., 2025), indicating that fast metabolic responses can be mediated at the metabolite level without requiring substantial changes in protein abundance. The stable proteome observed here is consistent with such a regulatory architecture, in which a constitutively maintained enzyme repertoire is modulated within minutes by metabolite pools, while transcriptional changes may adjust protein abundance only gradually across many cycles.

To further resolve the adaptive capacity of *Accumulibacter*, future studies will need to move beyond steady-state operation and deliberately perturb the system. Short-term challenges such as transient oxygen exposure during the anaerobic phase, abrupt shifts in carbon composition, or pulsed phosphate loading would test whether transcriptional regulation propagates to measurable protein-level changes when environmental predictability is disrupted. Pairing such experiments with time-resolved metabolomics would further show how metabolite pools and protein abundance jointly shape the response. Sustained stresses, including altered cycle lengths, reduced solids retention times, temperature shifts, or chronic nutrient imbalance, can impose selective pressures that may necessitate restructuring of the proteome itself.

The integrated multi-omics framework applied here indicates that regulatory dynamics in *Accumulibacter necessarius* are hierarchically organized, with extensive gene-level variability progressively buffered at higher functional levels and at the protein level. This organization offers a plausible explanation for how EBPR systems maintain stable performance under cyclic operation. More broadly, these results demonstrate how time-resolved multi-omics can be used to disentangle regulatory dynamics across molecular layers in engineered microbial ecosystems.

## Data Availability

Raw metagenomic sequencing data are available under BioProject PRJNA1537748, and raw metatranscriptomic sequencing data are available in the NCBI under BioProject PRJNA1298234. Genome-resolved datasets have been deposited in Zenodo under DOI 10.5281/zenodo.23180797, including the 1126 MAGs with GTDB taxonomy and CheckM quality information, the curated 53-MAG reference database used for downstream analyses. The metaproteomics raw data will be deposited in the PRIDE repository.

## Acknowledgement

We thank the Wastewater treatment Plant Thunersee (Thun, Switzerland) for providing access to the sludge. Many thanks to Stéphane Marquis, Alyssa Etter and Alessandro Scapuso (LBE, EPFL) for their help with reactor monitoring and molecular biology preparations, Emmanuelle Rohrbach for the molecular biology preparation work, Emmy Oppliger for the metaproteomic extraction and Lucas Hemmerle for his advice on metaproteomic analysis. Finally, we thank Sylvain Coudret (GR-CEL, EPFL) for the acetate and propionate quantification.

This work was supported by the National Center of Competence in Research (NCCR) Microbiomes financed by the Swiss National Science Foundation (grant number 180575). R. Pei receives funding from the European Union’s Horizon Europe research and innovation programme under the Marie Skłodowska-Curie grant agreement No. 101203185 (DOI: 10.3030/101203185).

