## Supplementary Information for "From dynamic transcripts to stable proteins: multi-omics revealed regulatory buffering of *Candidatus* Accumulibacter under cyclic conditions"

[2.] Proteomic Core Facility (PCF), Ecole Polytechnique Fédérale de Lausanne (EPFL), Lausanne, Switzerland

[#] Current Address: Division of Microbial Ecology, Centre for Microbiology and Environmental Systems Science, University of Vienna, Vienna, Austria
* These authors contributed equally
[+] Corresponding authors.

#### 1. Synthetic medium preparation

The synthetic medium was prepared from two concentrated stock solutions, a carbon solution (C) and a nitrogen–phosphorus solution (NP). The C stock contained 5.67 g/L sodium acetate trihydrate ($C_2H_3O_2Na \cdot 3H_2O$), 2.28 g/L sodium propionate ($C_3H_5O_2Na$), 0.889 g/L $MgSO_4 \cdot 7H_2O$, 2.2 g/L $MgCl_2 \cdot 6H_2O$, and 0.4 g/L $CaCl_2 \cdot H_2O$. The NP stock contained 1.671 g/L $K_2HPO_4$, 0.649 g/L $KH_2PO_4$, and 0.048 g/L allylthiourea ($C_4H_8N_2S$) to inhibit nitrification. In addition, 50 mL of trace-element solution was added per 10 L of NP stock. The trace-element solution contained 16.22 g/L $C_{10}H_{14}N_2Na_2O_8 \cdot H_2O$, 0.44 g/L $ZnSO_4 \cdot 7H_2O$, 1.012 g/L $MnCl_2 \cdot 4H_2O$, 7.049 g/L $(NH_4)_2Fe(SO_4)_2 \cdot 6H_2O$, 0.328 g/L $(NH_4)_6Mo_7O_{24} \cdot 4H_2O$, 0.315 g/L $CuSO_4 \cdot 5H_2O$, and 0.322 g/L $CoCl_2 \cdot 6H_2O$.

Both C and NP stock solutions were autoclaved. Before use, 250 mL of bicarbonate solution containing 0.933 g/L $NH_4HCO_3$ and 0.533 g/L $KHCO_3$ was added to the NP stock, which was then brought to a final volume of 10 L. For each reactor cycle, 120 mL of C stock and 120 mL of NP stock were mixed with 960 mL of distilled water and fed to the reactor, resulting in a final reactor COD concentration of 300 mg $O_2$/L.

#### 2. Amplicon Sequencing

Samples for 16S rRNA gene amplicon sequencing were collected weekly at the end of the anaerobic phase and processed as previously described (Adler & Holliger, 2020). Briefly, biomass was washed with 1x PBS, homogenized, and stored at -20 °C. DNA was extracted using the Maxwell automated DNA purification system (Promega) following lysozyme treatment (1 h, 37 °C). The V1-V2 region of the bacterial 16S rRNA gene was amplified using primers 27F/338R and Q5 High-Fidelity polymerase. Indexed libraries were prepared using the Nextera XT Index Kit v2, purified with AMPure XP beads, quantified by Qubit, pooled at 10 nM, and sequenced on an Illumina MiSeq v2 platform (2 x 250 bp) at the Lausanne Genomic Technologies Facility.

Reads were processed in DADA2 v1.32.0 (Callahan et al., 2016). Adapter sequences were removed using cutadapt v3.5 (Martin, 2011), followed by quality filtering and trimming (truncLen = c(225,225), maxEE = c(2,2), truncQ = 2, maxN = 0). Error models were learned separately for forward and reverse reads, sequences were dereplicated and inferred using pseudo-pooling, paired reads were merged, and chimeras were removed using the consensus method. Taxonomy was assigned using the MiDAS 5.3 database (Dueholm et al., 2022). ASVs were agglomerated to genus level using tax_glom in phyloseq v1.46.0, and genera reaching ≥3% relative abundance in at least one sample were retained for visualization (McMurdie & Holmes, 2013).

#### 3. Metagenomics

Processing of metagenomic reads was performed with the following bioinformatics tools with default settings unless specified otherwise. Raw metagenomic reads were first assessed for basic statistics using SeqKit v2.8.0 (Shen et al., 2024). Quality control was carried out with ATLAS v2.15.2 (Kieser et al., 2020), followed by normalization using BBNorm v39.06 (Bushnell, 2014), targeting an average sequencing depth of 320x. Reads with estimated depth below 2x were removed to improve assembly quality. Assembly of the normalized reads was performed using SPAdes v3.15.5 (Prjibelski et al., 2020) with the --meta flag. For comparison, normalized reads were compared with non-normalized reads assembled using MEGAHIT v1.2.9 (Li et al., 2015). Assembled contigs were post-processed with Anvi'o v8 (Eren et al., 2020) by simplifying contig names and discarding sequences shorter than 2500 bp.

Reads were then mapped to the assembled contigs using Bowtie2 v2.5.3 (Langmead & Salzberg, 2012), and binary alignment (BAM) files were generated using SAMtools v1.20 (Danecek et al., 2021). Coverage information was derived from both individual and merged BAM files to support genome binning. Binning was conducted using two non-competitive approaches, MetaBAT2 v2.12.1 (Kang et al., 2019) and SemiBin2 v2.1.0 (Pan et al., 2023), alongside the competitive approach VAMB v4.1.2 (Nissen et al., 2021). For competitive binning, additional reads from AGS metagenomes (Adler et al., 2022) were included to improve resolution.

Genome bins were refined using MAGScoT v1.0.0 (Rühlemann et al., 2022). The completeness and contamination of MAGs were evaluated using CheckM2 v1.0.1 (Chklovski et al., 2023), and taxonomic classification was assigned using GTDB-Tk v2.4.0 (release 220) (Chaumeil et al., 2022). As an optional step, the CAT/BAT toolkit v0.6.1 (Von Meijenfeldt et al., 2019) was used with the nr-euk database (04 April 2024) for additional taxonomic annotation. Dereplication of metagenome-assembled genomes (MAGs) was carried out using dRep v3.5.0 (Olm et al., 2017) with the following thresholds: ≥50% completeness, ≤10% contamination, and ≥98% average nucleotide identity for secondary clustering. CoverM v0.7.0 (Aroney et al., 2025) was used to estimate the relative abundance of MAGs and determine the fraction of metagenomic reads mapped to each genome. Finally, gene prediction was performed using Prodigal v2.6.3 (Hyatt et al., 2010) in metagenomic mode to extract protein-coding sequences and their corresponding amino acid translations. Gene annotation was performed using DRAM (Shaffer et al., 2020) and eggNOG using emapper-2.0.1 and emapper DB: 2.0 (Cantalapiedra et al., 2021).

#### 4. Metaproteomics

##### Sample preparation for mass spectrometry

Samples for metaproteomics were thawed on ice. For each extraction, 50 mg of biomass was resuspended in 300 µL lysis buffer (8M Urea) supplemented with a protease inhibitor cocktail (cOmplete Mini EDTA-free, Roche; one tablet per 10 mL of buffer). The samples then were sonicated with an amplitude of 60% for 5 cycles of 5 seconds. The sonicated sample was centrifuged at 13000 rpm at 4 °C for 15 minutes. The supernatant was recovered, snap-frozen in liquid nitrogen and stored at -80 °C. The extracted proteins were quantified using PierceTM BCA Protein Assay Kit (Thermo Fisher Scientific, USA).

20 µg of protein samples were loaded on a 12% SDS-PAGE gel and allowed for a short migration. The gel pieces containing the concentrated proteins were excised and washed twice with 50% ethanol in 50 mM ammonium bicarbonate (AB, Sigma-Aldrich) for 20 min and dried by vacuum centrifugation. Proteins were reduced with 10 mM dithioerythritol (Merck-Millipore) for 1 h at 56 °C followed by a washing-drying step as described above. Reduced proteins were alkylated with 55 mM iodoacetamide (Sigma-Aldrich) for 45 min at 37 °C in the dark followed by the same washing-drying step as described above. Proteins were digested overnight at 37 °C using mass spectrometry-grade Trypsin (Trypsin Gold, Promega) at a concentration of 12.5 ng/µl in 50 mM

AB supplemented with 10 mM $CaCl_2$. Resulting peptides were extracted in 70% ethanol, 5% formic acid (FA, Merck-Millipore) twice for 20 min, dried by vacuum centrifugation and desalted on SDB-RPS StageTips (Kulak et al., 2014).

**TMT Labelling**

For TMT labelling, cleaned and dried peptides were reconstituted in 12 µL of 100 mM HEPES pH=8 and 5 µL of TMT labels in solution (final concentration of TMT labels at 42 µg/µL in pure acetonitrile) was added. TMT Labelling was performed with the TMTpro 18-plex isobaric Mass Tagging Kit (Thermo Fisher Scientific) at room temperature for 90 min and reactions were quenched with hydroxylamine (final concentration of 0.4% (v/v)) for 10 min. Samples were distributed into two TMT sets. In each TMT set, an aliquot of a mix of all biological samples was included (bridge sample). The bridge samples allowed the results from the two TMT sets to be combined using the internal reference scaling (IRS) procedure described below. Each sample and bridge within a TMT set was labelled with a different labelling reagent. TMT-labelled samples were then pooled at a 1:1 ratio across all samples. The combined samples were then desalted using a 100 mg Sep-Pak C18 cartridge (Waters AG, Baden-Dättwil, Switzerland) and vacuum centrifuged. Pooled samples were fractionated into 12 fractions using an Agilent OFF-Gel 3100 system following the manufacturer's instructions. Resulting fractions were desalted on SDB-RPS StageTips (Kulak et al., 2014).

**Mass Spectrometry**

Each individual fraction was resuspended in 2% acetonitrile in 0.1% formic acid and nano-flow separations were performed on a Vanquish Neo nano UPLC system on-line connected with an Orbitrap Ascend Tribrid Mass Spectrometer (Thermo Fisher Scientific). A capillary pre-column (Acclaim Pepmap C18; 3 µm-100 Å; 2 cm x 75 µm ID) was used for sample trapping and cleaning. Analytical separations were performed at 250 nL/min over a 150 min biphasic gradient on an in-house packed capillary column (50 cm x 75 µm ID; ReproSil-Pur C18-AQ 1.9 µm silica beads; Dr. Maisch). Acquisitions were performed through Top Speed Data-Dependent acquisition mode using cycle time of 3 seconds. First MS scans were acquired over an m/z window of 400 to 1600 at a resolution of 120'000 (at 200 m/z). The most intense parent ions were selected and fragmented by high energy collision dissociation (HCD) with a normalized collision energy (NCE) of 35% using an isolation window of 0.7 m/z. Fragmented ion scans were acquired at a resolution of 45'000 (at 200 m/z) in the orbitrap with a Maximum Injection Time of 105 ms and selected ions were then excluded for the following 60 s.

**TMT Data Analysis**

Raw data were processed using Proteome Discoverer v.3.0. Spectra were searched with Sequest HT and MSFragger (Kong et al., 2017) against a custom protein database constructed from the predicted amino acid sequences of 53 MAGs recovered in the present study. Protein sequences from all MAGs were pooled and dereplicated using CD-HIT at 99% sequence identity, and the resulting database was supplemented with common MaxQuant contaminant entries. For peptide identification, enzyme specificity was set to trypsin and a minimum peptide length of six amino acids was required. Mass tolerance values were set to 10 ppm at parent level and 0.05 Da at fragment level. Carbamidomethylation (C), TMTpro tags (K residues and peptide N termini) were set as fixed modifications whereas oxidation (M) was considered as a variable one. Up to two missed cleavages were allowed and a 1% false discovery rate (FDR) cut-off was applied both at peptide and protein identification levels. A minimum of two peptides were required for protein identifications.

Resulting text files were processed through in-house written R scripts (v4.4.0). First a Sample Loading normalization (Plubell et al., 2017) was applied within each TMT set, assuming that the total protein amounts were equal across TMT channels. Briefly for each channel the reporter ion intensities were summed across all spectra, and the average across channels was calculated. Then, each channel was scaled to match its total intensity to the average. The multiplexing design of the experiment required a second normalization step to correct variations between the two TMT sets. IRS was here applied (Plubell et al., 2017) using the bridge samples. Assuming the protein concentrations are the same in the bridge samples from the two TMT sets, protein-specific scaling factors were calculated, based on their geometric means in the bridge samples. Additionally, a trimmed M-Mean normalization step was also applied using the package edgeR v4.0.14 (Robinson et al., 2010).

### Supplementary Figures

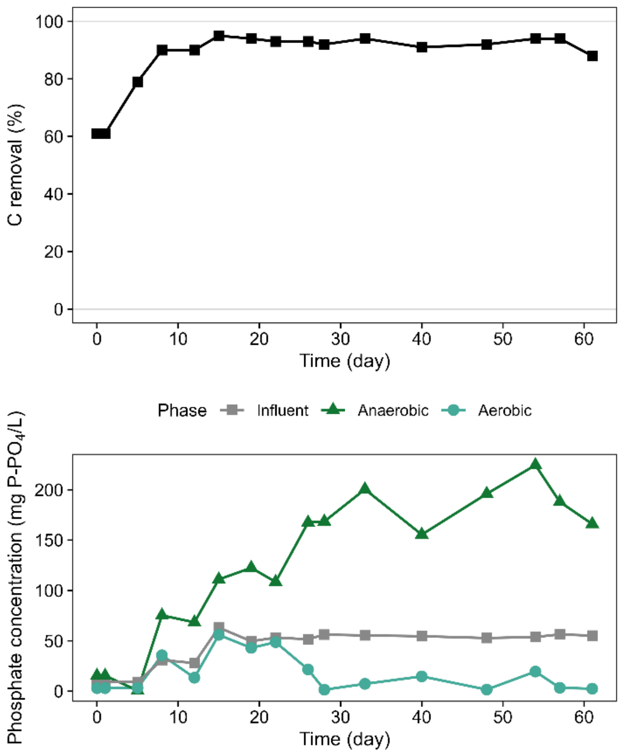


Figure S1. **Reactor performance during enrichment of the EBPR community. Top:** Carbon removal efficiency over time. **Bottom:** Phosphate concentrations measured in the influent, at the end of the anaerobic phase, and at the end of the aerobic phase.

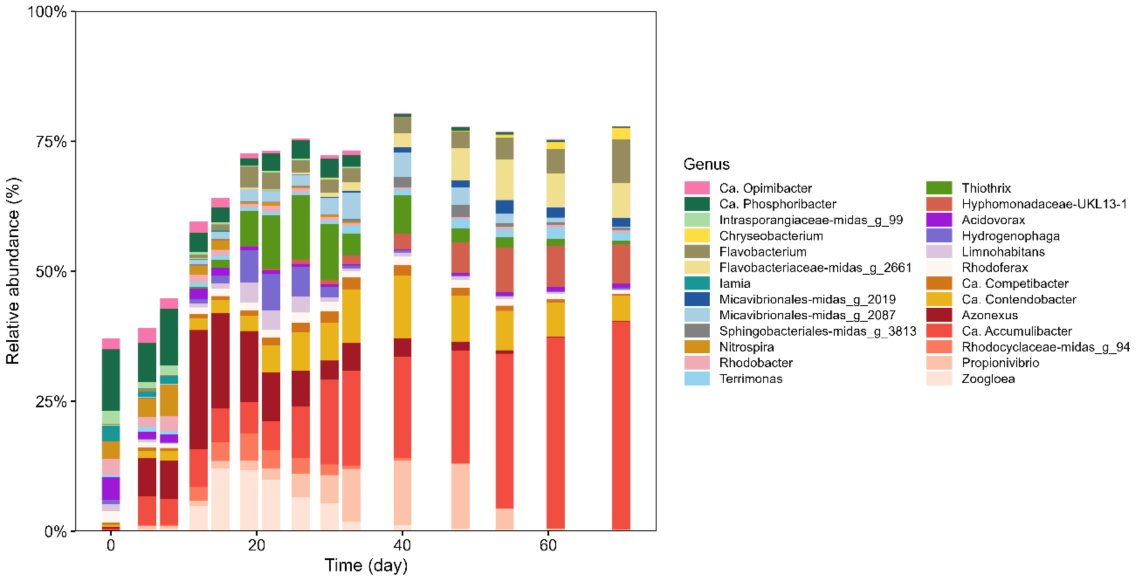


Figure S2. **Dynamics of the microbial community during reactor enrichment based on 16S rRNA gene amplicon sequencing.**

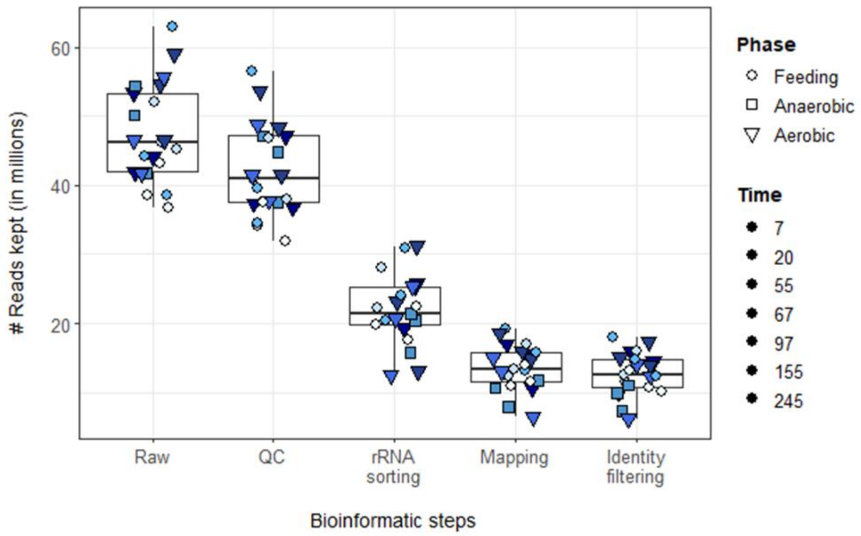


Figure S2. **Read retention across the metatranscriptomic bioinformatic workflow.**

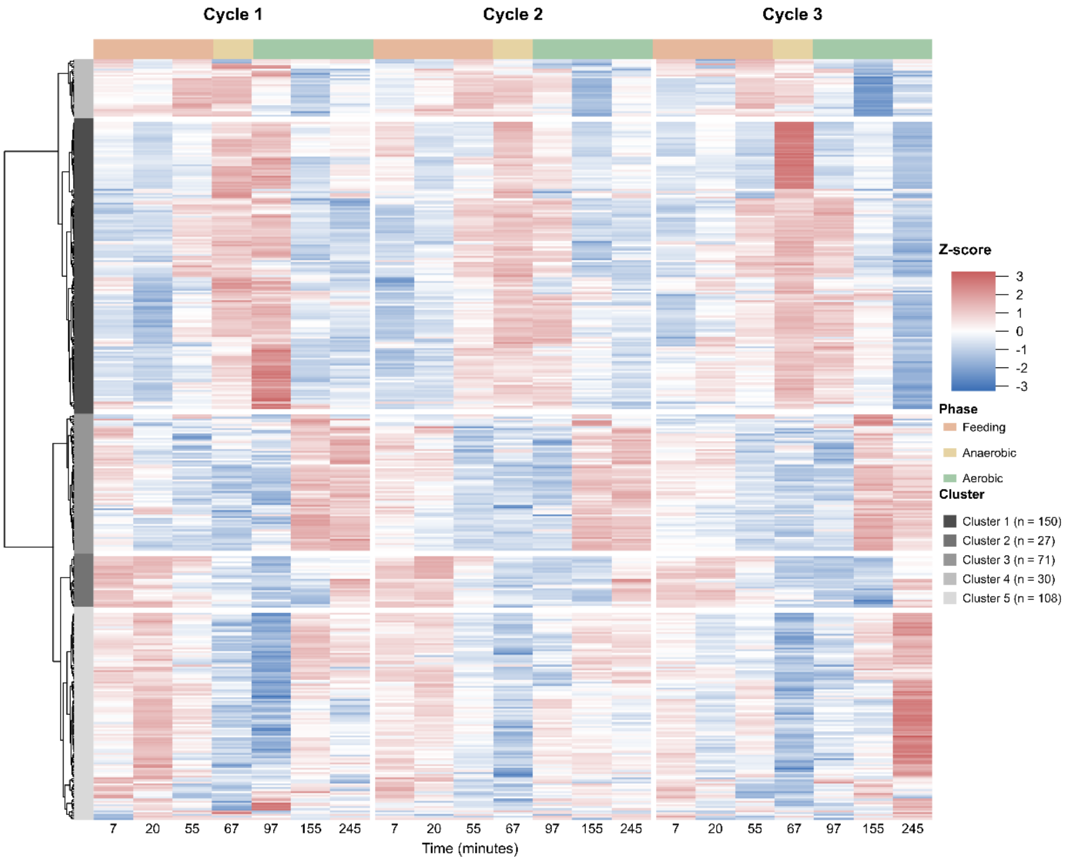


Figure S1. **Heatmap and hierarchical clustering of 386 differentially expressed genes across three consecutive EBPR cycles.** Rows represent individual DGE transcripts and columns represent the seven sampling times in each cycle. Expression values are shown as Z-scores, with red indicating higher and blue lower relative expression. Hierarchical clustering grouped the transcripts into five temporal expression clusters, with cluster sizes indicated in the legend.

#### Data Availability

Processed 16S rRNA gene amplicon sequencing reads after adapter removal and quality trimming are available in Zenodo under DOI 10.5281/zenodo.23180797.
